# SlytheRINs: using graph parameters and residue interaction networks to analyze protein dynamics and structural ensembles

**DOI:** 10.64898/2026.02.26.708270

**Authors:** Laura Shimohara Bradaschia, Matheus Epifane-de-Assunção, Marília V. A. de Almeida, Ândrea Kelly Ribeiro dos Santos, Umberto Laino Fulco, Ivanovitch Silva, Gustavo Antônio de Souza, Diego Marques Coelho, Gilderlanio Santana de Araújo, João Paulo Matos Santos Lima

**Author notes:** Corresponding author: Bioinformatics Multidisciplinary Environment (BioME), Digital Metropolis Institute (IMD), Universidade Federal do Rio Grande do Norte (UFRN), Av. Capitão-Mor Gouveia, S/N - Lagoa Nova, CEP: 59078-900, Natal, RN, Brazil.

## Abstract

Establishing the relationship between protein structure and functional behavior remains a significant challenge. The recognition that proteins are inherently dynamic, with functions often dependent on conformational changes, is increasingly accepted. Among computational approaches for elucidating protein properties, Residue Interaction Network (RIN) analysis has emerged as a powerful tool. However, conventional RIN analysis is constrained by its reliance on single, static protein structures, which fail to capture the flexibility inherent in dynamic protein folding transitions. To address these limitations, SlytheRINs is introduced as an interactive tool designed for comparative analysis of protein conformations via RINs. SlytheRINs enables dynamic ensemble analysis by decomposing interaction network data across multiple conformations of a single protein and providing detailed residue-interaction mapping across conformational changes via graph parameters in comparative plots. Applying these principles, the conformational variations of the wild-type and a pathogenic variant (G188R) of the human Glucose-6-Phosphatase (G6PC1) catalytic subunit were compared to identify fluctuations in both chemical interactions and graph features associated with conformational changes induced by the residue modification. The analyses identified key shifts in dynamic residue interactions in the protein variant that compromise substrate binding and the catalytic site, thereby elucidating the impact on G6PC1’s dynamic behavior and the resulting activity loss.

**Availability:** The source code for SlytheRINs is available on GitHub (https://github.com/evomol-lab/SlytheRINs), while the web tool and documentation are available at https://slytherins.streamlit.app.

## INTRODUCTION

Across all organisms, proteins are essential for sustaining life as they serve as the molecular instruments through which genetic information is expressed, ranging from structural components to catalytic functions in metabolic pathways (Caetano-Anollés et al., 2009; Hannah et al., 2023). The specific function that a protein assumes depends heavily on its three-dimensional conformation acquired during folding, which organizes the molecule’s primary structure into domains and catalytic sites (Dill et al., 2007). Consequently, studying protein structure yields relevant insights into physiological roles within cellular metabolism and sheds light on how genetic mutations may perturb function. However, the traditional paradigm of proteins as rigid, uniquely defined configurations had undergone a revolutionary shift in recent decades (Frixione and Ruiz-Zamarripa, 2023). Many proteins are now recognized as intrinsically disordered, dynamic, and pleomorphic structures that exhibit stochastic behavior rather than fixed conformations (Cui et al., 2025). This understanding has profound implications for structure-function relationships, as protein function fundamentally depends on conformational transitions among multiple states rather than on a single static structure (Bowman, 2024).

Chemical interactions between amino acids fundamentally determine protein folding and biological function (Newberry and Raines, 2019). To understand the importance of residue-residue interactions and their influence on protein function, Residue Interaction Network (RIN) analysis has emerged as a robust approach (Del Conte et al., 2024). Based on graph theory, these networks consist of nodes representing amino acid residues and edges representing adjacent physicochemical interactions. This method becomes even more valuable when applied to dynamic protein structures, such as conformational ensembles generated by molecular dynamics simulation (Begue et al., 2025).

Dynamic simulations and other conformation-generating methods, such as Normal modes and Elastic Network models, can produce hundreds to thousands of conformations, which are useful for mapping the multiple geometric states accessible to a protein, through which it may transition while performing its catalytic functions (Ma, 2005; Bahar et al., 2010). However, despite this advantage, the large conformational ensemble generated makes the comparative analysis of every residue interaction across networks derived from different proteins notably challenging. Consequently, the development of efficient computational methods is essential for analyzing large volumes of simulated data. To address this challenge, SlytheRINs was developed to perform comparative analysis of RINs derived from different protein structures using graph-based metrics and to summarize differences in physicochemical interactions for each residue. This tool facilitates the identification of crucial changes in the intra-protein network, enabling assessment of functional and structural changes resulting from sequence divergence between homologous proteins, polymorphisms, or mutations.

## FEATURES

SlytheRINs is an open-source, interactive Python application built with Streamlit (Streamlit Inc., 2026) that provides post-simulation network analysis using protein RINs. It uses the graph-based representation of chemical interactions to provide insights into residue-residue interactions across multiple frames comprising a simulation trajectory or conformational ensemble. The tool integrates two main tabs: (i) Comparative RIN Analysis, (ii) Chemical Interaction Analysis, each with distinct data analysis and visualization characteristics (Figure 1).

**Figure 1.**
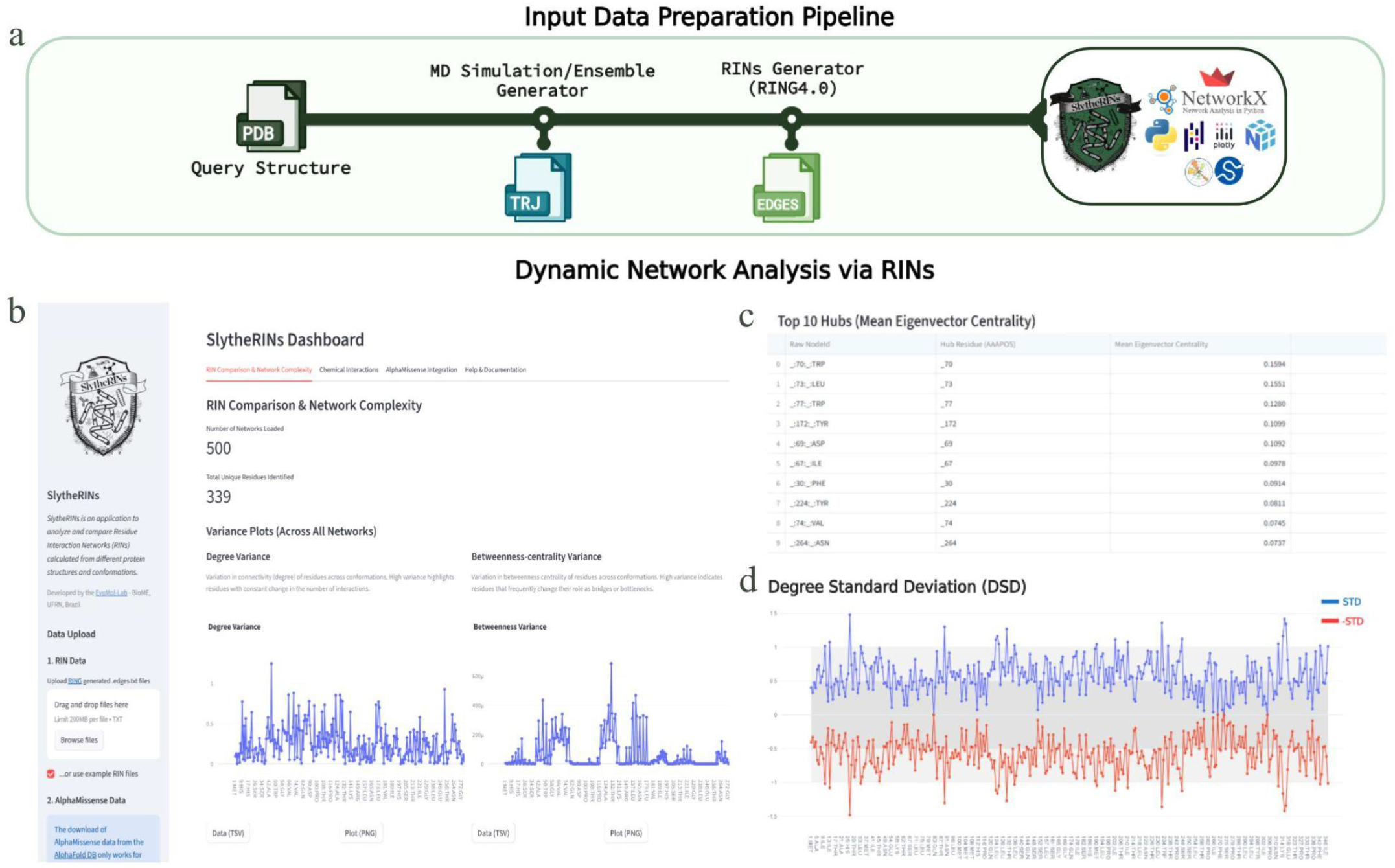
**(A)** General SlytheRINs input data preparation pipeline. **(B)** User Interface in Comparative RIN Analysis Tab, followed by **(C)** Top 10 Hubs described by Eigenvector Centrality and **(D)** Degree Standard Deviation Plot.

As its focus is on multiple conformational data analysis, the primary input to SlytheRINs consists of multiple.edges.txt files describing the edges of RINs for protein structures. Those files must be pre-generated by applying RIN-builder tools, such as RING 4.0 (Del Conte et al., 2024), to molecular dynamics trajectory or conformational ensembles files of interest (usually a structure file with several model coordinates). This process builds a distinct network for each trajectory frame, enabling further comparison of residue-residue interactions across conformations (Figure 1a).

The Comparative RIN Analysis in SlytheRINs comprises dynamic network analysis to capture residue interaction patterns and topological variation across multiple conformations. This is achieved through two main analytical approaches: the computation of residue-level centrality descriptors and global network properties for each individual conformation, and the statistical analysis of topological descriptors at both the residue and network levels to characterize variation across the conformational ensemble.

To compose the backend of the first analytical process, *networkX* (Hagberg et al., 2008) is used to assess network topology and compute essential centrality measures for each conformation (frame or model). Then, the second analytical process characterizes residue connectivity fluctuations and network organization across the conformational ensemble by performing statistical analyses of the obtained measures, summarized in Table 1. For network topology, the tool calculates assortativity and also performs two statistical tests on the variation of degree values per residue. The first one, a paired T-test, tests whether a residue’s degree (across networks) differs significantly from the average degree of all other residues in those same networks. The second one tests whether the variation in each residue’s degree values is significantly different from what would be expected under a random (Poisson) process. These analyses allow the user to identify crucial residue sites associated with structural and conformational variation.

Upon completion of the analytical process, the tool interface presents summary metrics, including the number of networks loaded and the number of amino acid residues (nodes). Subsequently, variance plots of degree, betweenness centrality, and clustering coefficient for each residue in each topology are provided (Figure 1b). The interface also includes mean and standard deviation plots for triangle counts, degree, and eigenvector centrality (hubs), as well as a table listing the top 10 residue hubs based on their mean eigenvector centrality values (Figure 1c). SlytheRINs further reports each residue’s Degree Standard Deviation (DSD) (Fonseca et al. 2020) (Figure 1d), which quantifies the variability in the number of interactions for a residue. Low DSD values indicate stable connections, while high DSD values suggest dynamic behavior. The Comparative RIN Analysis tab also includes additional analyses of network complexity, both per network and across all networks, to help identify potential power-law distributions characteristic of scale-free networks.

The second tab, Chemical Interaction Analysis, quantifies and plots the mean counts and standard deviations for different types of chemical interactions (e.g., HBOND, VDW, IONIC, PIPISTACK) for each residue across all networks. Chemical interaction labels are extracted from the “Interaction” column of the input edge files, using the same attributes as in RIN construction. SlytheRINs adopts the nomenclature defined by RING4.0 (Del Conte et al., 2024). To facilitate user engagement, all modules include examples and sample data, enabling execution of all described steps and processes and familiarization with the tool’s outputs and their interpretation. Additionally, users can generate a consolidated PDF report or download individual plots and data in.tsv,.csv, and.png formats.

## USAGE EXAMPLE

### Assessment and comparison of the conformational variations of the wild-type and a pathogenic variant (G188R) of the human Glucose-6-Phosphatase (G6PC1) catalytic subunit using SlytheRINs

The performance of SlytheRINs was evaluated by comparing the wild-type human protein G6PC1 with a missense variant at position 188, which substitutes Glycine (G) for Arginine (R). G6PC1, also known as G6Pase-α, is a small integral membrane protein (357 residues) embedded in the endoplasmic reticulum (ER) membrane. It functions as an indispensable subunit of human Glucose-6-phosphatase (G6Pase), which hydrolyzes glucose-6-phosphate (G6P) in the ER lumen. The Glycine at position 188 is located in an intramembrane helix (Fig 2A and 2B), and the Gly188Arg variant (rs80356482) is associated with a complete loss of activity in Glycogen Storage Disease Type Ia (GSDIa).

**Figure 2.**
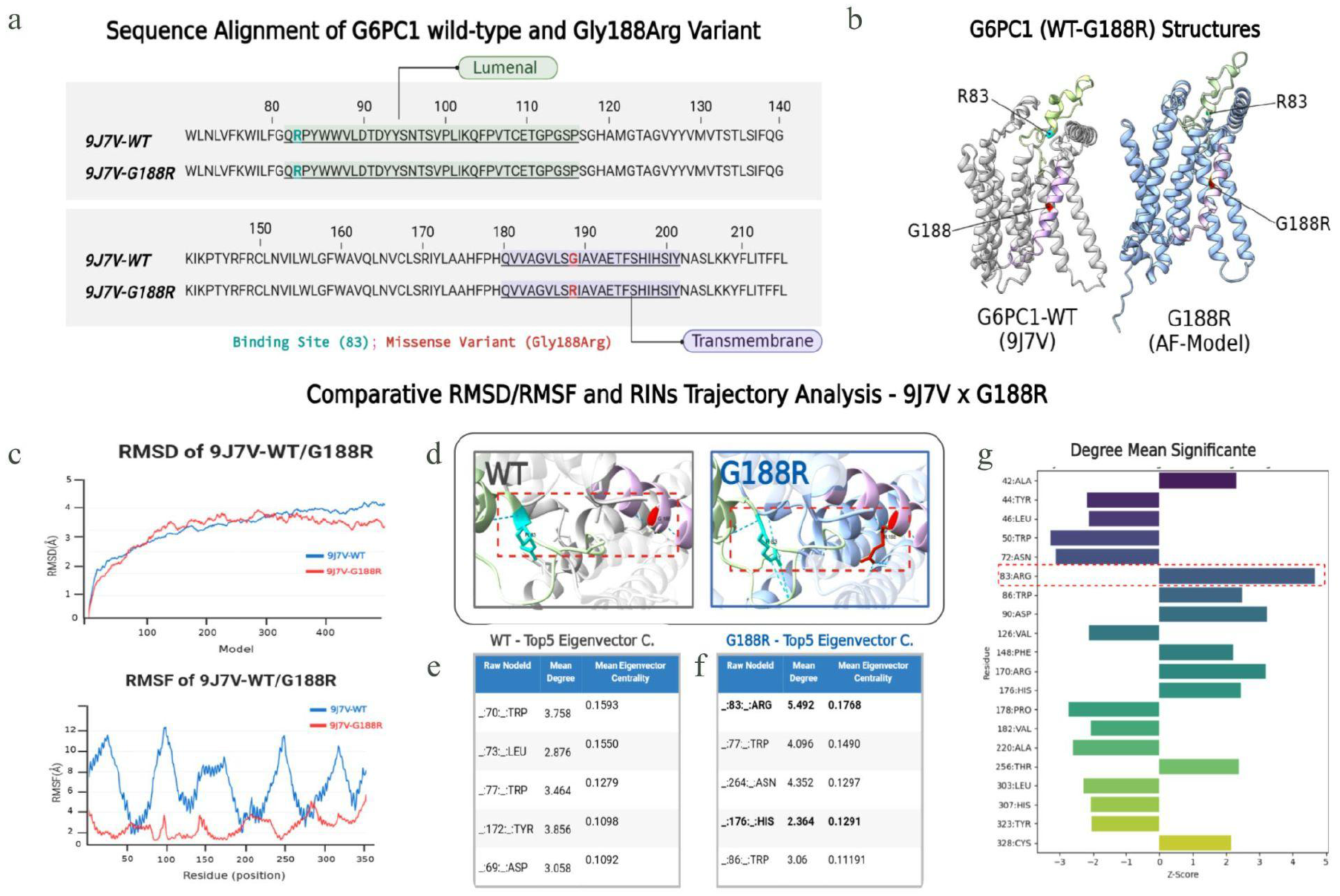
**(A)** Sequence alignment between the native G6PC1 sequence retrieved from 9J7V PDB structure against Gly188Arg variant, binding site (R83-Cyan), and missense variant (G188-Red) locations are highlighted, as well as each topological domain. (**B)** 9J7V-WT (white) and G188R (blue) structures. Both the R83, G188 location and its topological domain are highlighted in the same colors as the sequence in 1A. **(C)** Root Mean Square Deviation (RMSD) to monitor and compare overall structure changes over time, and Root Mean Square Fluctuation (RMSF), to identify residue flexibility changes between proteins from 9J7V-WT (blue) and G188R (red). **(D)** and **(E)** SlytheRINs top5 Eigenvector Centrality and **(F)** Positional Zoom Visualization for 9J7V-WT and G188R. **(G)** Z-Score standard deviation by residue position between 9J7V-WT and G188R Degree Means. R83 and H176 (G6PC1 binding and active sites, respectively) positions show significant changes in connectivity, with the greatest deviation in the degree mean between the two structures.

### Data Acquisition and Processing

The Apo and native state G6PC1 structures were retrieved from the PDB (ID: 9J7V), while the G188R variant structure was modeled using AlphaFold3 (Abramson et al. 2024) based on the sequence retrieved from the UniProtKB/SwissProt Variant page (VAR_005246). Trajectories for both molecules were simulated using the NMSIM web server (Krüger et al. 2012) with a normal mode, coarse-grained approach. RINs for each trajectory were calculated using RING 4.0 (Conte et al. 2024), which generated edge output files that served as input to SlytheRINs.

### Comparative Analysis

Both the PDB wild-type (9J7V) and the G188R variant structures were analyzed separately using the SlytheRINs web tool, with results exported in.tsv,.png, and report formats for each. Complementary RMSD and RMSF plots were generated for each trajectory to support the network analysis. Structural and sequence visualizations were produced using ChimeraX (Meng et al. 2023) and PyMOL (Schrödinger, LLC. The PyMOL Molecular Graphics System, Version 2.4.0 https://www.pymol.org/).

RMSD and RMSF calculations indicated a shift in residue fluctuation in the G188R variant, demonstrating considerably less flexibility compared to the wild-type 9J7V-WT (Fig. 2C). Structural visualization (Fig. 2D) highlights the emergence of new interactions originating from position 188 that are absent in the native state. Analysis using the Comparative RIN Analysis module of SlytheRINs identified a significant difference in eigenvector centrality between the two molecules (Fig. 2E-2F). Notably, residues R83 and H176 emerged as the most influential nodes within the G188R structure. Assessment of the difference in mean degree/connectivity further confirmed R83 as a highly connected residue in the variant structure (Fig. 2G).

Biologically, Arginine at position 83 is one of the two known binding sites of G6PC1, and H176 is an active site essential for the protein’s catalytic activity. By integrating RMSD, RMSF, and biomolecular insights with RIN analysis across conformations, SlytheRINs enabled visualization of the distal impact of the G188R mutation. Although the mutation is not located directly at the active site, it significantly alters the network topology of critical catalytic and binding sites (R83 and H176).

## FINAL REMARKS AND CONCLUSION

Conventional RIN analysis tools rely on single, static protein structures, whereas SlytheRINs addresses the critical gap in analyzing dynamic protein ensembles by treating conformational variants as interconnected networks rather than isolated snapshots. By enabling comparative analysis across hundreds to thousands of conformations from molecular dynamics simulations, the web tool quantifies residue interaction stability and variability across conformational landscapes, providing information that would be practically unfeasible to extract manually. This approach aligns with the current understanding that protein function fundamentally depends on conformational transitions between multiple states rather than a single rigid structure (Bowman, 2024; Cui et al., 2025).

Comparative analysis of the native G6PC1 structure and the pathogenic G188R variant demonstrates how SlytheRINs analyses can reveal distal mutation effects through changes in network topology. Although the mutation is located in an intramembrane helix distant from the active site, SlytheRINs identified significant alterations in eigenvector centrality and degree connectivity at R83 and H176, the known binding and catalytic sites, respectively (Xia et al., 2025). By integrating RMSD, RMSF data, and network metrics, the tool enabled visualization of how the G188R substitution propagates its effects through the residue interaction network, ultimately disrupting critical functional sites and leading to complete loss of enzyme activity in GSDIa (Hannah et al., 2023). This demonstrates the utility of comparative RIN analysis for understanding the impacts of mutations beyond direct structural perturbations.

The open-source nature and web-based accessibility of SlytheRINs democratize sophisticated network analysis for researchers without extensive computational expertise. As molecular dynamics simulation data and conformational analyses continue to grow, tools that efficiently analyze ensemble properties and export comprehensive reports become increasingly essential. Thus, SlytheRINs provides a framework for understanding how local interaction changes contribute to global functional consequences in both fundamental research and clinical variant interpretation.

## Acknowledgements

The authors are indebted to the High-Performance Computing Center (NPAD) at UFRN for providing computational resources.

## Artificial Intelligence Disclosure

The authors used generative AI technologies for code auditing and performance optimization (Gemini® and GitHub-CoPilot®) and for stylistic editing of the manuscript (Grammarly®). Throughout this process, the authors maintained full control over the research design and interpretation of results; the AI acted solely as a technical and linguistic aid.

